# Accelerating Monte Carlo modeling of structured-light based diffuse optical imaging via “photon sharing”

**DOI:** 10.1101/2020.02.16.951590

**Authors:** Shijie Yan, Ruoyang Yao, Xavier Intes, Qianqian Fang

## Abstract

The increasing use of spatially-modulated imaging and single-pixel detection techniques demands computationally efficient methods for light transport modeling. Herein, we report an easy-to-implement yet significantly more efficient Monte Carlo (MC) method for simultaneously simulating spatially modulated illumination and detection patterns accurately in 3-D complex domains. We have implemented this accelerated algorithm, named “photon sharing”, in our open-source MC simulators, reporting 13.6× and 5.5× speedups in mesh- and voxel-based MC benchmarks, respectively. In addition, the proposed algorithm is readily used for accelerating the solving of inverse problems in spatially-modulated imaging systems by building Jacobians of all illumination-detection pattern pairs concurrently, resulting in a 12.4-fold speed improvement.

https://doi.org/10.1101/2020.02.16.951590

## 1. INTRODUCTION

Diffuse optical tomography (DOT) and fluorescence molecular tomography (FMT) with near-infrared light have impacted an increasing number of clinical and pre-clinical applications over the last decade, such as monitoring breast tumor response to therapy and quantifying drug delivery in small animals [1]. The appeal of these model-based imaging methods is largely elicited by their non-ionizing nature, high sensitivity, functional imaging, and multiplexing capabilities, in addition to relatively low cost. Recently, new DOT/FMT instrumentation methodologies based on structured-light, including spatial-frequency domain imaging (SFDI), have been proposed to improve upon the traditional point-source illumination schemes [2, 3]. Enabled by spatial light modulators (SLMs) such as digital micro-mirror devices (DMDs), the illumination/detected light can be spread/acquired over large surfaces. This allows the total illumination power to be significantly higher than those in point-source modalities without damaging the tissue being probed. As a result, the signal-to-noise ratio (SNR) of the collected measurements is significantly enhanced, especially when performing wide-field patterned detection, leading to dramatically reduced acquisition times. Moreover, modulations via patterns can be imparted both in the illumination and detection optical paths to enable 3D quantitative probing over a large field-of-view [4, 5]. However, quantitative and efficient analyses of wide-field spatially-encoded imaging systems impose new challenges for light transport modeling and image reconstruction techniques.

Among methods used to model light propagation in turbid bio-tissues, the Monte Carlo (MC) method is recognized as the gold standard thanks to its accuracy and generality over a wide range of conditions, including the mesoscopic regime [6], low albedo tissues, and early-arriving photons [7]. Benefiting from the rapid development in parallel computing hardware, and especially that of graphical processing units (GPUs), MC has become the forward model of choice in many DOT and FMT applications. A typical MC simulation involving millions of photons can now be accomplished within seconds on a personal computer (PC), making it viable to apply such accurate forward models in more challenging DOT/FMT data processing applications where repeated simulations are required. Additionally, MC provides high flexibility to accommodate the implementations of complex illumination and detection strategies, making it a convenient platform for exploring and optimizing structured-light based imaging systems. To meet this growing need, we have previously developed open-source structured-light-capable modeling platforms – Monte Carlo eXtreme (MCX) [8] and Mesh-based Monte Carlo (MMC) [9,10].

These open-source simulators are able to accurately and efficiently simulate complex illumination patterns and flexible photon detection approaches, making them well suited for the development of structured-light imaging systems [9,11]. However, solving the inverse problems for wide-field spatially encoded DOT/FMT experiments requires the processing of hundreds of illumination and detection pattern combinations [5], strongly necessitating additional speed accelerations to achieve practical feasibility. To mitigate this computational burden, Belanger et al. [4] proposed simulating a pencil beam source and convolving the solution with each illumination pattern to produce all fluence distributions required to compute the whole Jacobian. However, this approach works only for homogeneous media with presumed translational invariance in space and time [12]. In the case of SFDI systems with Fourier patterns, Gardner et al. derived a complex photon weight from the SFD radiative transfer equation (RTE) to incorporate the effects of spatial modulation on a photon’s initial weight [13], allowing solutions for multiple spatial frequencies to be estimated in a single MC simulation. However, this approach cannot be used with other encodings, such as Hadamard, Haar or speckle patterns [11].

In this letter, we report a general approach to performing MC simulations for structured light illumination and detection strategies with the ability to obtain the full set of forward solutions for all illumination or detection patterns using only a single forward simulation. This feature greatly reduces the computational burden of structured light transport modeling by leveraging our understanding that a photon’s trajectory in an MC simulation is independent of the weight imposed by source or detection patterns. As a result, the propagation of a photon can be effectively “shared” between all structural patterns. We refer to this approach as “photon sharing”.

The remainder of the paper is organized as follows. In Section 2, we first describe this approach in the context of forward modeling using a sample single-pixel wide-field DOT platform [4, 5]. Then we demonstrate the benefit of leveraging this approach to rapidly compute the perturbed measurements and sensitivity matrix, i.e. Jacobian, for inverse problems. In addition, the considerations on memory optimization to implement this algorithm on the GPU are discussed. In Section 3, we validate the proposed algorithm and quantify speed improvements compared to conventional approaches. The speedup ratios are obtained for a set of commonly used structured-light patterns. Finally, we summarize our findings and discuss future works.

## 2. MATERIALS AND METHODS

### A. Paralleled forward model for multiple light patterns

To better describe the proposed algorithm, we show a sample schematic of a single-pixel-camera imaging system [4,5] in Fig. 1, where structured illumination is combined with single-pixel detection to provide wide-field spatially-modulated DOT. Modulated by the illumination DMD, a set of structured light patterns are sequentially projected to the surface of the tissue to be imaged. The diffuse light that transmits through the phantom is encoded with another set of patterns generated by the detection DMD and redirected to a photon detector, such as a photomultiplier tube (PMT). For each of the patterns, the spatial intensity variance is represented using a two-dimensional (2-D) raster image, *I_s_* and *I_d_*, with a normalized intensity value (floating-point numbers between 0 and 1) assigned at each pattern pixel. Without losing generality, in Fig. 1, we assume *N_s_* illumination patterns and *N_d_* detection patterns are needed.

**Fig. 1.**
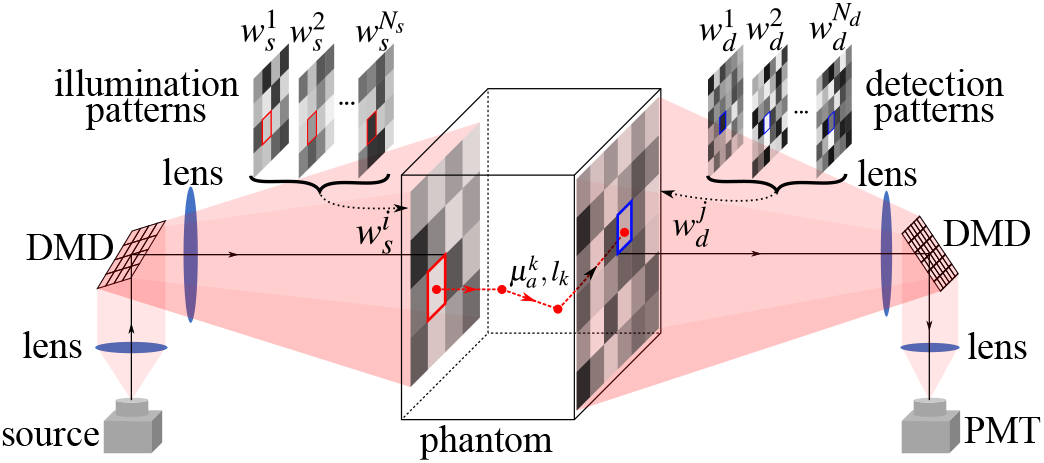
An illustration of a structured-light based diffuse optical imaging system with single-pixel camera detection.

In sequential MC simulations, when simulating the t-th source pattern, a photon packet’s initial weight is first assigned as the normalized source pattern intensity value, 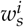, at the launch position, indicated by a red-square in Fig. 1. After propagating the photon packet over its trajectory, consisting of a discrete set of line segments with lengths *l_k_* and absorption coefficients 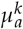, indicated by red-arrows in Fig. 1, the photon exits the domain at a location, indicated by a blue-square, within the field- of-the-view of the detector. In conventional MC simulations, this results in an additional multiplication by the normalized detector pattern weights, 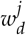 for the *j*-th detector pattern.

In our accelerated simulations, we recognize that the time-consuming photon trajectory simulations within turbid media are independent of the assigned initial weight 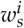 and the detector weight 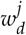; therefore, they can be effectively “shared” among all source and detector patterns. To enable this parallel simulation, we first assign a uniform value 1.0 for all launched photon packets. In the meantime, the initial weights corresponding to all source patterns (see red-squares in Fig. 1) are concatenated into a vector 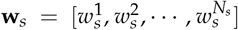 and stored in the local memory of the computing thread that handles such photons. As the photon traverses across different voxels or elements of the domain, the weight loss, Δ*w*, is accumulated in voxels to generate the volumetric fluence, i.e. the forward solution. At the *k*-th path segment, the weight losses for all source patterns can be computed simultaneously as

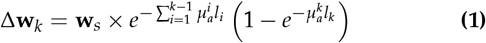

and accumulated in the memory for all source patterns after each propagation step *k*. Therefore, after a single MC forward simulation, we can obtain the fluence maps for all sources.

### B. Accelerated perturbation Monte Carlo formulation for single-pixel imaging system

The perturbation MC method (pMC) [14] is often used to rapidly recompute detector readings without rerunning the full MC simulation. Here we propose an accelerated pMC approach to quickly obtain measurements for all combinations of patterned sources and detectors using just a single MC forward simulation.

The rasterized source patterns (*I_s_*) and detector patterns (*I_d_*) are pre-defined and thus, instead of storing the initial weight vector, **w**_*s*_, for all detected photons, we only store the linear index, *M*, by treating the *i*-th source pattern 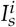 as a 1-D vector. In other words, with the saved launch position index M, we can recreate the initial weight vector by 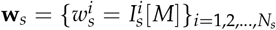. When a patterned source is used in conjunction with a nonpatterned detector (such as a disk), one can rapidly recompute the detected photon weight *W* using the stored index *M* and the partial pathlength vector {*L_t_*}_*t*=1,2,…,*N*_*t*__, i.e. the accumulative path-lengths for each tissue type, for *N_t_* tissue types, as

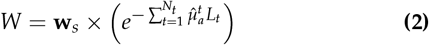

where 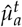 denotes the perturbed absorption coefficient for the *t*-th tissue type.

The above formula can be extended for patterned detectors. In this case, if a photon is detected by a patterned detector, we can also store the exit-position (or detector pattern index), with which one can compute the detector weight vectors 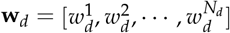. By performing an outer product between the initial weight vector **w**_*s*_ and **w**_*d*_ for a given detected photon, we can recreate its detected weights *w_i,j_* at all source/detector pattern pairs as

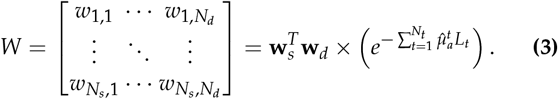

### C. Accelerated Jacobian computation for structured light DOT

#### C.1. The adjoint Monte Carlo

The Jacobian formulations derived in [15] based on the adjoint Monte Carlo (aMC) can be readily applied to the dual-DMD system shown in Fig. 1. For the combination of the *i*-th (*i* = 1,2, …, *N_s_*) illumination and *j*-th (*j* = 1,2, …, *N_d_*) detection pattern, referred to as the pattern pair (*i, j*), the continuous-wave Jacobians for absorption (*μ_α_*) and scattering (*μ_s_*) perturbations can be expressed as

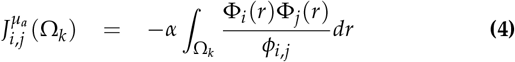

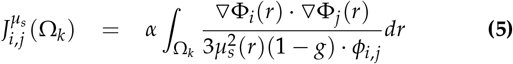

where Ω_*j*_ denotes the *k*-th spatial region; *α* denotes the scaling factor between diffuse reflectance and fluence at the boundary [15]; Φ_*i*_ (*r*) and Φ_*j*_ (*r*) are the normalized fluence distributions for the *i*-th source pattern and *j*-th detection pattern, respectively; *ϕ_i,j_* is the measurement of the total diffuse reflectance or transmittance (*R*) for pattern pair (*i, j*) computed by

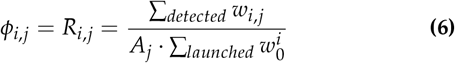

where 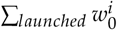 is the total simulated photon weights for the *i*-th source pattern; *Σ_detected_ W_i,j_* is the total detected photon weights for pattern pair (*i, j*), with *w_i,j_* computed by Eq. 3; *A_j_* is the effective area covered by the *j*-th detection pattern. By applying the proposed algorithm, building Jacobians via aMC only needs two MC simulations: one forward simulation of all illumination patterns to compute Φ_*i*_ (*r*) and *ϕ_i,j_*, and another one for Φ_*j*_ (*r*) by projecting detection patterns as the time reverse “adjoint” sources.

#### C.2. The “replay” Monte Carlo

In [15], we reported a direct approach, referred to as photon “replay”, to rapidly create the Jacobians derived from pMC [16]. Briefly, the photon replay algorithm involves two steps – first, in the “baseline” MC simulation, the random number generator (RNG) seeds of all detected photons are stored; secondly, in the replay step, detected photons are relaunched using the RNG seed saved earlier. The detected photon weights are precomputed before the re-propagation, during which the weighted photon path lengths and weighted scattering numbers are deposited to produce spatially and temporally resolved Jacobians. By applying Eq. 3 for computing the detected photon weights during the output storage, we can then apply Eqs. (6–7) in Ref [15] to obtain the Jacobians using photon replay.

### D. Memory optimization

Please note that the storage of initial weight vectors **w**_*s*_ within each computing thread may produce substantial computational overhead if handled improperly. To minimize such overhead, we strongly recommend reordering the source/detector pattern array dimensions so that the inner-most (i.e. the fastest) index is the dimension corresponding to the pattern index. Such a memory layout allows a thread to access all source/detector pattern weights over a contiguous section of memory, which is often more efficient than accessing non-contiguous locations.

The above suggested memory layout is particularly important on memory-limited processors such as GPUs. Loading/saving data for all patterns within a GPU thread can be done much more efficiently when the pattern-related data are stored along contiguous memory spaces. In such cases, the GPU can access data with significantly less latency. As a matter of fact, on modern GPUs, every global-memory read retrieves a contiguous 128-byte cache line [17], corresponding to 32 single-precision numbers. This suggests that reading one or 32 single-precision values from the global memory has roughly the same memory latency. Therefore, storing pattern data using a pattern-index-first order can maximize such cache efficiency.

In addition, as we showed in Eq. 1, the **w**_*s*_ vector is repeatedly used for every step along a photon’s trajectory. To minimize the memory latency in our GPU implementation [8], we store **w**_*s*_ using the high-speed shared memory [17].

## 3. RESULTS AND DISCUSSIONS

In this section, we first demonstrate the improved speed of MCX and MMC, in runtime (s) per pattern, by comparing the proposed algorithm against the conventional approach where simulations of structured light patterns are performed sequentially. In Fig. 2(b), the speedup ratio is further quantified by testing over varying numbers of simultaneously simulated patterns (or groups) for a fixed total pattern count (128). Group size of 1, 2, 4, 8,16, 32, 64 and 128 are tested. All patterns have the same dimension of 32 × 32 pixels. For MCX [8], the patterns are projected to a 40 × 40 mm^2^ illumination area through the bottom surface of a 60 × 60 × 30 mm^3^ slab-shaped homogeneous optical phantom, with an absorption coefficient *μ_a_* = 0.01 mm^-1^, scattering coefficient *μ_s_* = 10 mm^-1^, anisotropy *g =* 0.9, and refractive index n = 1.37. A heterogeneous mesh model generated from the Digimouse atlas is used for benchmarks on MMC (sample images shown in Fig. 2(a)). The mesh data and the optical properties are detailed in [18]. The illumination area is a 20 × 40 mm^2^ rectangle on the back of the mouse. In addition, the influence of pattern complexity on speed is assessed for a fixed group size (32). Light patterns with varying resolutions are tested, ranging from 8 × 8 to 1024 × 1024 [see Fig. 2(c)]. A total of 10^9^ and 10^8^ photons are launched for MCX and MMC simulations, respectively. All benchmarks are performed on a desktop running Ubuntu 16.04, with an NVIDIA TITAN V GPU for MCX and an AMD Ryzen 9 3950X CPU for MMC.

**Fig. 2.**
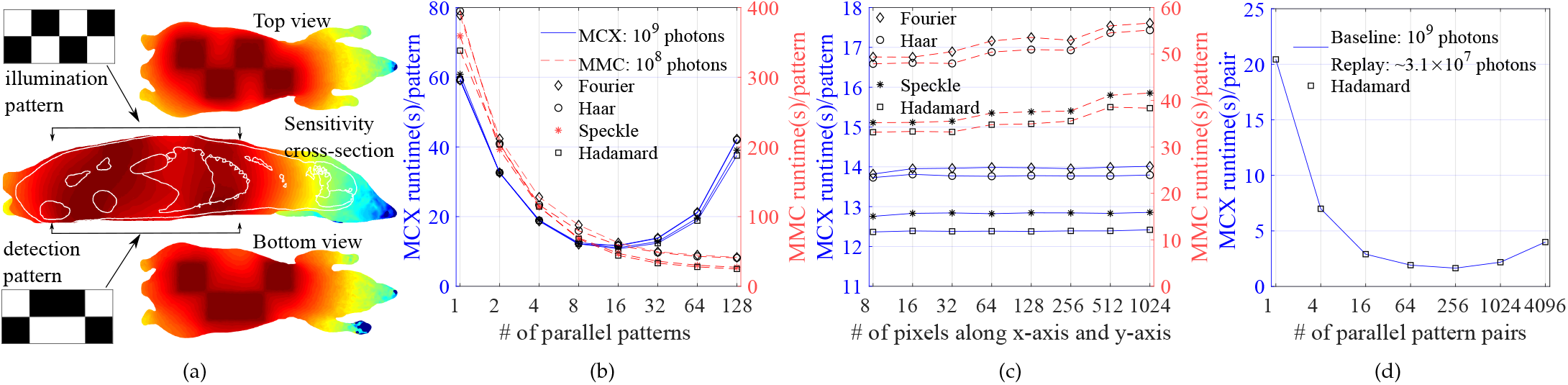
Sample (a) fluence (top/bottom) and sensitivity (middle) maps in a heterogeneous mouse atlas and speed assessments by (b) varying the number of simultaneously simulated pattern count (for a total of 128 patterns), (c) varying isotropic pattern pixel dimensions, and (d) varying the simultaneously replayed pattern pairs for a Jacobian with 64 source and 64 detector patterns.

In all benchmarks, the fluence distributions produced by the proposed algorithm show excellent agreement with those computed individually through conventional approaches (not shown). In Fig. 2(b), with increased pattern numbers, the runtime per pattern for MMC decreases monotonically, achieving a speedup ratio ranging between 9.38× (Fourier, 389.0 vs. 41.5 s/pattern) and 13.6× (Hadamard, 338.2 vs. 24.9 s/pattern) when simulating all 128 patterns together. However, on GPU-based MCX, simulating 16 patterns per batch produces the highest speedup – ranging between 5.03× for Fourier (59.2 vs. 11.7 s/pattern) and 5.5× for Hadamard (59.5 vs. 10.7 s/pattern). The performance degradation when simulating more than 16 patterns together on the GPU is likely a result of reduced active thread-blocks due to increased use of shared memory [17].

In Fig. 2(c), the average runtime over a range of pattern resolutions are reported for four pattern bases, including Hadamard, Haar, Fourier and speckle (all patterns are normalized between 0 – 1). From this plot, we can observe that the speed is relatively steady, indicating that this algorithm can readily support high resolution light patterns with low computational overhead. In addition, we also observe slight speed variations between different simulated patterns. The speed is roughly inversely correlated to the fill factor of the patterns (i.e. percentage of non-zero values) due to the associated memory overhead. For instance, Fourier patterns lead to the slowest runtimes because there are very few zeros, while Hadamard patterns show the highest speed as they are binary patterns with around 50% zeros.

Next, we quantify the performance improvement of photon “replay” for MCX. The simulation setup is similar to the MCX benchmark discussed earlier except that Hadamard patterns are used not only for illumination, but also to cover a 40 × 40 mm^2^ area of the top surface to perform single-pixel detection, as shown in Fig. 1. In Fig. 2(d), the average replay runtime against varying numbers of total illumination-detection pattern pairs is reported. The highest speedup ratio is achieved at 12.4× (20.4 vs. 1.7 s/pair) when the sensitivity matrices of 256 pattern pairs are computed simultaneously. The slight speed decrease when using more than 256 simultaneous pattern pairs may also be a result of limited shared memory space.

## 4. CONCLUSION

In summary, we reported a general and highly efficient algorithm to accelerate Monte Carlo simulations in emerging structured-light and single-pixel detection based imaging experiments. Compared to conventional sequential approaches, this algorithm is capable of simulating multiple source patterns simultaneously. We show that such acceleration can be extended to single-pixel camera systems with detector patterns and generalized for pMC and sensitivity calculations.

Enabled by the proposed algorithm, the CPU-based MMC demonstrated a monotonic increase in simulation speed with increasing pattern numbers and achieved 9.38 × to 13.6 × speedups in benchmarks of a Digimouse atlas. In addition, GPU-based MCX achieved the highest speedup of over 5-fold at groupings of every 16 patterns. Further increasing the simultaneous pattern number showed a drop in speed on the GPU due to the impact of shared memory utility. Finally, we combine “photon sharing” with our “photon replay” [15] algorithm, resulting in a 12.4 × overall speed improvement. This improved MC algorithm is specifically optimized for structured-light based diffuse optical imaging techniques and can be a valuable tool towards the development of fast, quantitative imaging platforms, paving the way for bedside studies of dynamic physiology and/or fluorescence guided interventions. The photon sharing feature has been incorporated into both of our MC simulators and can be freely downloaded from http://mcx.space/.

## 5. ACKNOWLEDGMENT

This work is supported by the National Institute of Health under grants R01-GM114365, R01-EB026998, and R01-CA204443 (QF) and R01-EB019443, R01-CA207725, R01-CA237267, R01-CA250636 (XI).

